# Mothers face immediate, but family-size dependent, costs of sons in preindustrial Finland

**DOI:** 10.64898/2026.04.08.717249

**Authors:** Euan A Young, Leon van Dorp, Mirkka Lahdenperä, Virpi Lummaa, Hannah L Dugdale

## Abstract

The expensive son hypothesis posits that mothers incur higher fitness costs when caring for sons versus daughters in species with male-biased size dimorphism. Evidence for maternal survival costs of sons in humans is limited to shortened overall lifespans; whether having more sons reduces short-term survival during reproductive years is unknown. Here, we utilised life-history data from 5,456 mothers from preindustrial Finland to examine whether mothers with more sons had reduced survival within one year of their last birth. While mothers with few children but more sons showed no differences in survival, at higher family sizes, mothers with more sons had increasingly lower survival. These differences peaked at ∼0.4% lower survival per son among mothers with five children, suggesting accumulated physiological costs of sons. These differences then declined and reversed among mothers with more children, potentially due to selective disappearance of frailer mothers. Our results suggest that studies focusing on post-menopausal mothers may bias estimates of the fitness costs of sons and reproductive costs more broadly. We recommend future research further examines the overlooked short-term fitness costs of sons during reproductive years, which is vital for understanding how life-history trade-offs, sexual dimorphism, and their interaction have shaped human evolution.

## INTRODUCTION

According to life-history theory, individuals suffer fitness constraints with increased allocation to reproduction (Kirkwood, 1977; Stearns, 1992). The expensive son hypothesis is an extension of this framework, positing that mothers incur greater fitness costs when bearing and raising sons rather than daughters in species with male-biased size dimorphism (Clutton-Brock, 1991). Increased fitness costs to mothers bearing sons have been widely found in such species and include, for example, reduced future reproductive success (Weiss et al., 2023) and survival [(Froy et al., 2016), reviewed in (Invernizzi, Lemaître, et al., 2024)].

In humans, like many mammals, sons are on average larger than daughters (Lindenfors et al., 2007), meaning sons demand ∼10% more energy intake for mothers during pregnancy (Tamimi, 2003) and ∼0.05 kg more breast milk per day (Da Costa et al., 2010). Many studies have examined the fitness costs of sons for human mothers (Invernizzi, Lemaître, et al., 2024), but our understanding is limited in two ways. First, studies have focused on long-term fitness costs on survival [e.g., sons reducing post-menopausal lifespan (Helle et al., 2002)] with short-term fitness costs being limited to reproductive factors [e.g., sons delaying future reproduction (Mace & Sear, 1997)]. Second, studies focusing on survival have almost entirely focused on mothers surviving to menopause [but see (Grandi et al., 2023; Helle & Lummaa, 2013)] with mixed results (Invernizzi, Lemaître, et al., 2024). This focus is despite there being no reason that these costs should only manifest after menopause (Clutton-Brock, 1991; Helle & Lummaa, 2013), and evidence that sons increase the chances of caesareans (Eogan, 2003): a medical intervention which responds to life-threatening complications for the mother during labour. If sons impact the survival of mothers before menopause, studies focusing only on mothers reaching menopause will (1) underestimate fitness costs of sons (Helle, 2017) and (2) may bias estimates if mothers surviving to menopause are, for example, biased towards less frail individuals (Doblhammer & Oeppen, 2003). Understanding whether sons reduce mothers’ short-term survival during their reproductive years is thus crucial.

Here, we utilise historical life-history data from 5,456 mothers from preindustrial Finland to examine whether mothers had reduced survival up to one year after their last birth if: (1) their last birth was a son or (2) if they had, in total, birthed relatively more sons than daughters. In our study population, during this period, both family sizes and maternal mortality are higher than in industrialized populations and some evidence of short-term costs of sons has already been accrued: mothers have delayed and reduced chances of reproduction after having male versus female co-twins (Lummaa, 2001); and reduced overall post-reproductive survival with more sons, with the associated mortality risk being highest in the period soonest after the last birth and declining thereafter (Helle & Lummaa, 2013). We also tested whether a mother’s total number of children and socioeconomic status moderated the effect of having more sons on survival. We predicted that mothers from lower socioeconomic statuses and/or who have had more children would show increased fitness costs of sons (Hurt et al., 2006).

## METHODS

### Study population

We used life-history records of mothers across rural Finland who had their last birth during the preindustrial period, before 1890 (N = 8,140). The data were collected from parish birth, marriage, death, and religious service attendance records kept by the Lutheran church, allowing individual life-histories to be constructed even in the case of migration. Industrialisation in Finland began slowly in the late 19^th^ century, with manufacturing still only employing 20% of the population as late as 1910 (Voutilainen, 2016). While vaccines against smallpox became available from 1880 onwards (Ukonaho et al., 2022), at the end of the 19th century, infectious diseases remained among the most common causes of death (Saarivirta et al., 2012). Overall, in preindustrial Finland, the standard of living was low: the median lifespan was 24, and ∼40% of children would die before their 15^th^ birthday. The mating system was also strictly monogamous; both divorce and extra-marital affairs were outlawed (Sundin, 1992).

### Selected data

We restricted our sample to mothers whose last birth was after 1749 (n = 7,928), after which the church was required by law to record all births, movements, marriages, and deaths in the country (Luther, 1993). We also included only mothers who had their reproductive records complete (as determined by genealogists; n = 6,581), and recorded mothers’ ages at last birth, whether their last birth was a twin or triplet (i.e., a multiple birth, yes or no), and the sex and birthdates of all children. We recorded whether a mother survived at least one year after her last birth to encompass the entire span of pregnancy-associated deaths (Tikkanen et al., 2020). For this, we used death records and, for mothers without death records (18%), classified mothers as survivors if they were documented in our records more than one year after their last birth (e.g., attending a religious event). Consequently, we removed a further 13 mothers who had appeared in the records after her last birth but within one year because survival after one year was unknowable. We also focused on mothers who had married once to remove heterogeneity in individual life histories (n: excluded = 597), recorded the region the mother lived in (n: Central Finland = 1,887, Eastern = 434, Northern = 1,446, or Southwestern = 1,689, missing = 258), and assigned an occupation derived socioeconomic status, which was the highest known status from either the mother or her husband (n: lower = 1,145, intermediate = 1,962, upper = 2,349, missing = 203), leaving 5,456 mothers (Figure S1).

### Statistical analysis

For these mothers, we modelled their short-term survival using Bayesian generalised linear mixed models (GLMMs) using a logit link function in *R* 4.3.2 (R Core Team, 2023) with *brms* 2.22.0 (Bürkner, 2018) and the Markov chain Monte Carlo sampler *Rstan* 2.32.7 (Stan Development Team, 2025). We examined the costs of sons by including a categorical variable of the sex of the last birth (male or female, and in the case of multiple births, male if at least one child was male) and, as a numeric variable, the proportion of all of a mother’s children who were sons. In models we controlled for: as continuous variables, a mother’s total number of children and age at last birth, adding squared terms due to potential non-linearity of trade-offs (Cohen et al., 2020); and, as categorical variables, the multiple birth status of the last birth (i.e., singletons or multiple birth [twins/triplets]), the region, and socioeconomic status. We also tested for a three-way interaction and all two-way interactions between the proportion of children who were sons, the total number of children, and socioeconomic status. To control for variation among birth years in maternal survival, we accounted for the year of the mother’s last birth as a random effect. We also accounted for variation among families in maternal survival by including a family random effect, but kept this as a supplementary analysis because many mothers did not have a recorded parent (n missing = 2,210), leading to a large reduction in the sample size when among-family variation was accounted for (N = 3,246). Mothers with recorded parents had, on average, 1.41 sisters also recorded as mothers in the data (range 1-5).

We evaluated evidence for our hypotheses using the Probability of direction (Pd) parameter. Pd values <0.95 were considered as non-significant, 0.95-0.975 as marginally significant, and >0.975 as significant. Non-significant interactions and squared terms (not included in an interaction) were removed, least significant first. We ran four chains for all models, for 12,000 iterations, sampling every 10 iterations, after burning the first 2,000, giving 4,000 posterior draws. Convergence was checked by confirming all Rhat parameters were approximately 1 and by visually checking that chains resembled “hairy caterpillars”. We also performed posterior predictive checks using the *pp_check()* function in *brms*. Default uninformative priors were used: fixed effects, flat, and random effects, a student’s t-distribution. Median posterior estimates and 95% credible intervals are shown: in tables, for fixed and random effects; and, in the text, for contrasts which show the predicted difference in survival between different values of variables from the models. Model predictions were made using posterior samples, using intermediate socioeconomic status, baseline predictors for other categorical variables, and median values for continuous variables, unless otherwise stated. Contrasts where 95% credible intervals do not overlap zero are equivalent to Pds > 0.975.

## RESULTS

Of all mothers having their last birth between 1750 and 1889 (N= 5,456), 8% died within the year following this birth (n = 446), and half of these were within the first 42 days [i.e., post-natal (WHO, 2010), Figure S2]. 52% of last births were to sons (n = 2,822), and mothers had a median of five children (range = 1-17), with 95% having 10 or fewer. Considering all children, mothers had a slightly son-biased sex-ratio (proportion of children who were sons = 51%), but the sex ratio varied considerably between mothers (range = 0-100%). This variation reduced at higher parities while remaining slightly biased towards sons: for example, among mothers with 5 children (n = 590), 16% had one or fewer sons and 22% had four or more; and among mothers with 10 children (n = 206), 4% had two or fewer, while 7% had eight or more (Figure S3).

We found no evidence for fitness costs of sons on maternal survival considering only the sex of their last birth (Pd < 0.975, Table 1). In fact, albeit statistically non-significant, mothers whose last birth were sons had higher survival during the year following birth than those with daughters (predicted survival difference = 0.5% [95% CIs, -0.4 - 1.4%]). However, considering their full reproductive histories revealed that mothers with more sons had similar, lower, or higher survival depending on the total number of children the mothers had (interaction between number of children and proportion of children who were sons, Pd > 0.975, Figure 1, Table 1). For example, among mothers who only had one child, mothers had similar survival if they had one son versus one daughter (predicted survival difference = -0.5% [-2.1% – 1.2%]). However, among mothers with four children, mothers having only sons had a lower survival probability than those having only daughters (predicted survival difference = -2.1% [-4.3% – -0.3%]) which also persisted for mothers with median numbers of children (i.e., five, -2.1% [-4.6% – 0.0%]). These reductions in survival for mothers having a greater proportion of sons then began to reverse among mothers with more children, such that amongst mothers with 10 children, mothers having only sons had higher survival than those having only daughters (albeit the difference was statistically non-significant, predicted survival difference = 12.6% [-2.1% – 33.3%]). The differences in survival of mothers with more sons became even larger amongst mothers with even more children, but with a high degree of uncertainty, owing to few mothers having more than 10 children (n = 222, Figure S4).

**Table 1:** Generalised linear mixed model results showing conditional associations between predictors and maternal survival one year after last birth using the full preindustrial Finland dataset (N = 5,456). Estimates and variances reported are the posterior distribution median and 95% credible intervals, rounded to three significant digits. Significant effects (Pd, probability of direction > 0.975) are in bold. Model results with non-significant interactions included are shown in Table S1.

| Fixed effect | Estimate [95% CI] | Pd |
| --- | --- | --- |
| Intercept | 1.24 [-1.28 – 3.68] | 0.837 |
| Age at last reproduction | -0.0589 [-0.217 – 0.103] | 0.758 |
| **Age at last reproduction^2** | **0.00338 [0.000908 – 0.00582]** | **0.998** |
| Sex (male) | -0.149 [-0.427 – 0.127] | 0.845 |
| Number of children | -0.0645 [-0.391 – 0.257] | 0.654 |
| Number of children^2 | -0.0101 [-0.0421 – 0.0233] | 0.728 |
| Proportion of children who were sons | 0.294 [-0.495 – 1.11] | 0.766 |
| **Socioeconomic status (middle)** | **-0.902 [-1.56 – -0.258]** | **0.998** |
| Socioeconomic status (lower) | 0.152 [-0.594 – 0.918] | 0.642 |
| **Multiple birth (yes)** | **-0.706 [-1.17 – -0.176]** | **0.994** |
| Region (eastern Finland) | -0.195 [-0.565 – 0.202] | 0.832 |
| **Region (northern Finland)** | **-0.318 [-0.579 – -0.0471]** | **0.989** |
| Region (south-west Finland) | 0.00823 [-0.256 – 0.265] | 0.524 |
| **Number of children x Proportion of children who were sons** | **-0.517 [-1.02 – -0.044]** | **0.981** |
| **Number of children<sup>2</sup> x Proportion of children who were sons** | **0.0647 [0.00967 – 0.119]** | **0.991** |
| **Number of children x Socioeconomic status (middle)** | **0.484 [0.194 – 0.782]** | **1** |
| **Number of children<sup>2</sup> x Socioeconomic status (middle)** | **-0.0472 [-0.0732 – -0.0214]** | **1** |
| Number of children x Socioeconomic status (lower) | 0.0673 [-0.329 – 0.445] | 0.634 |
| Number of children<sup>2</sup> x Socioeconomic status (lower) | -0.0105 [-0.0477 – 0.0316] | 0.683 |
| **Random Effects** | **Variance** | **Levels** |
| Birth year of last child | 0.445 [0.29 – 0.613] | 138 |

**Figure 1:**
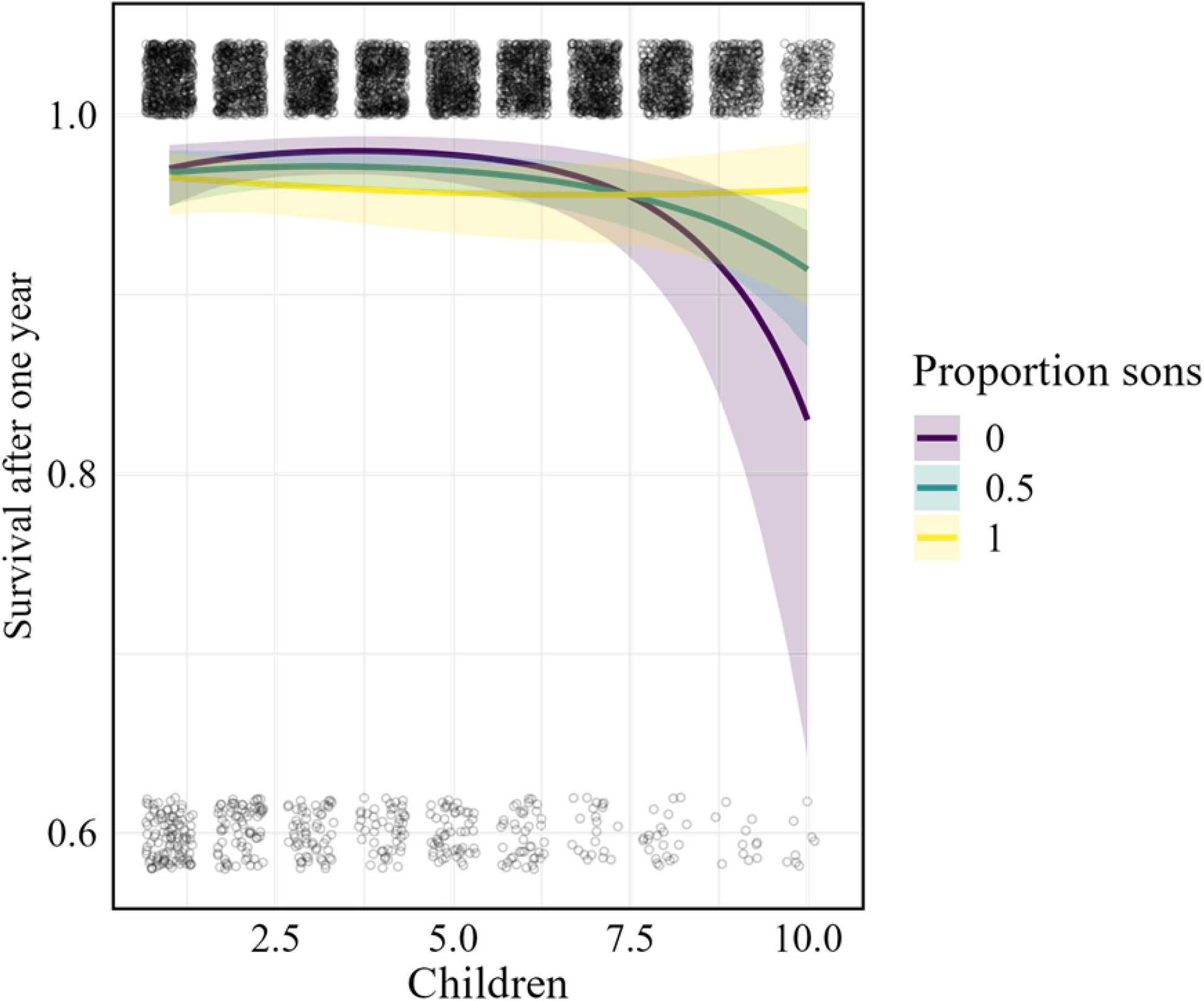
Predicted survival probabilities of mothers according to their number of children and the proportion of those children who were sons from a generalised linear mixed model performed on the full dataset (N = 5,456), from preindustrial Finland. For visualisation, we predict for mothers who had only daughters (purple), an equal proportion of sons and daughters (turquoise), and only sons (yellow), and from the 5^th^ to 95^th^ percentiles for the number of children (1-10). Predicted survival probabilities extended across all numbers of children are plotted in Figure S4. Survival probabilities were predicted using posterior samples and median values or baseline values for other predictors: age at last reproduction, 38.7; Central Finland; sex of last child, male; intermediate socioeconomic status; and singleton births. Shaded areas denote 95% credible intervals. Points show surviving (above), and deceased mothers (below) are jittered for visualisation. Full model results are shown in Table 1.

To understand the extent to which these differences in survival across mothers with more sons were driven by the differences in survival after one year among mothers with very large family sizes, we ran a supplementary analysis of only mothers with 10 or fewer children (95th percentile for number of children, N = 5,234). Here, the differences in survival between mothers with different sex ratios and numbers of children were marginally significant, suggesting that this result was partially driven by the survival difference among mothers with more than 10 children (Pds = 0.963-0.973, Table S2 and S3). However, the overall patterns of survival between mothers with more sons across different numbers of children remained consistent (Figure S5, Table S3): among mothers with four children, mothers having only sons had lower survival than those having only daughters (predicted survival difference = -2.9% [-5.7% – - 0.3%]) and, among mothers with 10 children, mothers having only sons had a higher survival probability than those having only daughters (albeit the difference was again statistically non-significant, predicted survival difference = 7.1% [-3.3%, 28.4%]). The differences in survival for mothers with more sons across different numbers of children were also robust to a supplementary analysis accounting for among-family variation in survival (N = 3,246, Figure S6, Pds > 0.975, Table S5, among-family variation = 1.27 [0.439 – 1.98]).

Forty-three percent of mothers belonged to the higher socioeconomic status group (n = 2,349 upper; 36%, n = 1,962 intermediate, 21%, n = 1,145 lower) and the differences in survival across mothers with different proportions of sons and total children were consistent across mothers from all socioeconomic statuses (interaction between socioeconomic statuses, number of children, and proportion of children who were sons, Pd < 0.975, Table S1). We found some evidence that mothers differed in survival between socioeconomic statuses across mothers with different numbers of children (Table 1, Figure S7), but this association appeared to be driven only by mothers of intermediate socioeconomic status having lower survival among mothers with 10 or more children and this association entirely disappeared in our supplementary analysis of only mothers with 10 or fewer children (Tables S2-4, Figure S8). In fact, mothers had very similar survival within one year of last birth across socioeconomic groups (predicted survival upper socioeconomic status = 95.8% [94.3% – 96.9%], intermediate = 95.7% [94.2% – 96.9%], and lower = 96.7% [95.3% – 97.7%], Table S3).

These results accounted for differences in survival between mothers with different ages at last birth and last birth years, from different geographical regions and families, and whose last birth was a singleton or multiple birth. Mothers had their last child between ages 15 and 51 (median = 39) with the survival of mothers being higher for mothers with later ages at last birth: mothers with their last birth at age 39 had a survival chance of 97.2% [95.7% – 97.9], with mothers having their last birth after age 46 having at least a 99.4% [99.0% – 99.7%] chance of surviving for one year after their last birth (Pd > 0.975, Table 1, Figure S9). Mothers from northern Finland had lower survival than those from central and southwestern (predicted survival difference both = -1.1% [-2.3% – -0.2%]) but not eastern Finland (predicted survival difference = -0.5% [-1.9% – 1.0%], Table 1, Pd > 0.975). Finally, mothers whose last birth was a multiple birth (twins or triplets) had lower survival (predicted survival difference = -2.9% [-0.6% – 6.3%], Pd > 0.975, Table 1), and a mother’s survival also varied across families and birth years (Tables 1 and S5).

## DISCUSSION

Here, we investigated, for the first time, the short-term survival costs of sons on reproductive-age mothers using data from preindustrial Finland and found that the costs of sons differed among mothers with different numbers of children. While mothers with one child had a similar survival within one year of birth regardless of whether they had a son or not, mothers showed costs at higher numbers of children, such that among mothers with five children (the median for this population), mothers with more sons had reduced survival probabilities of ∼0.4% per son (Figure 1). These results support the expensive sons hypothesis (Clutton-Brock, 1991) and shed new light on short-term costs among reproductive-age mothers. However, contrary to predictions, these costs then steadily weakened and reversed among mothers with even more children. We discuss potential explanations and key implications below.

We found no survival costs of sons when considering only the sex of the child at last birth, and finding these costs present only when considering several births suggests an accumulation of physiological costs rather than mortality due to one-off effects, such as more birth complications purely due to the larger body sizes of sons. Instead, these accumulated physiological costs would appear to mediate a mother’s health and manifest as an increased risk of mortality in the year following birth, and particularly in the first four weeks (Figure S2). Previous studies examining long-term survival costs of sons have suggested a causal role of testosterone (Helle et al., 2002) – which has an immunosuppressive function for mothers (Foo et al., 2017) – or increased inflammation (Galbarczyk et al., 2021) in explaining these effects. Whether such mechanisms could influence maternal mortality within a year of birth is unknown, and underlying mechanisms are difficult to unravel within our historical study system. Our results do not contradict the existence of long-term costs; in fact, they align closely with previous work on this population, which found that survival risks of having sons were highest for mothers in the period closest after births and declined thereafter (Helle & Lummaa, 2013). However, our results emphasize that ignoring short-term survival costs paid by reproductive-age mothers may mean underestimating the fitness costs of sons (as argued previously in the context of reproductive costs across both sexes, Helle, 2017) and may partially explain the mixed evidence in favour of survival costs of sons in humans (Invernizzi, Lemaître, et al., 2024). Although we cannot disentangle these competing mechanisms in our study, we hope to inspire future studies that may elucidate the mechanisms underlying the short-term survival costs of sons, which have been overlooked in humans.

Contrary to our prediction (Hurt et al., 2006; Stearns, 1992), short-term survival costs of sons then diminished among mothers with more than five children, and, among mothers with eight or more children, more sons were instead associated (non-significantly) with higher maternal survival (Figure 1). The statistical significance of these associations was strengthened by the inclusion of mothers with extremely high family sizes (i.e., more than 10 children), but the patterns and magnitude of these associations were very consistent across all models (Figure S5), even when accounting for among-family differences in survival (Figure S6). One reason for the reversal of these costs could be the selective disappearance of frailer individuals among mothers having many sons, which biases survival estimates in demographic studies of reproductive costs (Doblhammer & Oeppen, 2003). Selective disappearance could drive the reversal of survival differences among mothers with more sons if frailer mothers are less likely to survive bearing sons, leading to a more robust selection of mothers among mothers with both many children and a higher proportion of sons than daughters. This aligns with our observation that mothers with more sons had higher survival among mothers where selective disappearance is strongest (i.e., mothers with higher family sizes), and a recent study showing that mothers with the poorest health (approximated through infant mortality) showed the greatest costs of sons (Invernizzi, Bergeron, et al., 2024). Alternatively, sons may be playing an active role in increasing maternal health so that mothers with many sons have reduced costs at later births, such as by increasing farm production, as previously speculated in this population (Nitsch et al., 2013). However, the evidence for such an ability of sons is weak, and we believe the selective disappearance of frailer mothers is a more likely explanation.

These results have important implications not only for the expensive sons hypothesis, but also for studies looking at the costs of reproduction in humans more broadly. Our results suggest that, in addition to underestimating the fitness costs of sons by excluding mortality during reproductive years, studies focusing on mothers surviving to menopause may further bias estimates because they examine reproductive costs on a more robust subset of mothers that have already survived to menopause. The effect of frailty or selection bias in the cost of reproduction literature has been known for decades (Doblhammer & Oeppen, 2003), yet studies continue to exclude mothers dying before reaching menopause (e.g., Young et al., 2025). Excluding mothers dying before age at menopause can be considered an example of collider bias, where the exposure variable (number of children or sons) and response variable (survival) both affect the likelihood of being selected in a study (survival to age at menopause; (Munafò et al., 2018; Tönnies et al., 2022). This effect is further evidenced in our study by mothers showing improved survival with later ages at reproduction, a correlate of their total number of children. Moreover, as far as we are aware, the only previous study on the survival costs of sons among reproductive-age mothers from preindustrial populations also found evidence for fitness costs of sons (Helle & Lummaa, 2013), while studies focusing only on post-menopausal mothers report mixed evidence (Invernizzi, Lemaître, et al., 2024). Future studies of the expensive sons hypothesis and costs of reproduction would therefore benefit from the inclusion of mothers dying during reproductive years, or explicit modelling of frailty differences among post-menopausal mothers.

In conclusion, our results support the application of the expensive sons hypothesis to humans (Clutton-Brock, 1991) and suggest that studies may underestimate these costs when ignoring the short-term survival costs of mothers before menopause. While these costs may be considered small (∼0.4% lower survival within a year of birth per son among mothers with median family sizes), it is worth emphasising that these effects were detectable within purely-observational variation in the numbers of children, sex ratios, and mortality among mothers in our preindustrial population. This adds to the evidence of evolutionary constraints on the human family size in the form of maternal mortality (Gagnon, 2015; Hurt et al., 2006), which, when occurring during reproductive years, has particularly severe fitness consequences. Nevertheless, quantifying the fitness costs of sons on reproductive-age mothers is complex, and these results require replication. We suggest future studies model non-linear (e.g., quadratic) survival costs and test for dependencies of reproductive costs across mothers to better understand the role of life-history trade-offs, sexual dimorphism, and their interaction in human evolution.

## ACKNOWLEDGEMENTS

We thank E Postma and the Dugdale Research Group for feedback throughout the project.

