## Supplementary materials for "Mothers face immediate, but family-size dependent, costs of sons in preindustrial Finland"

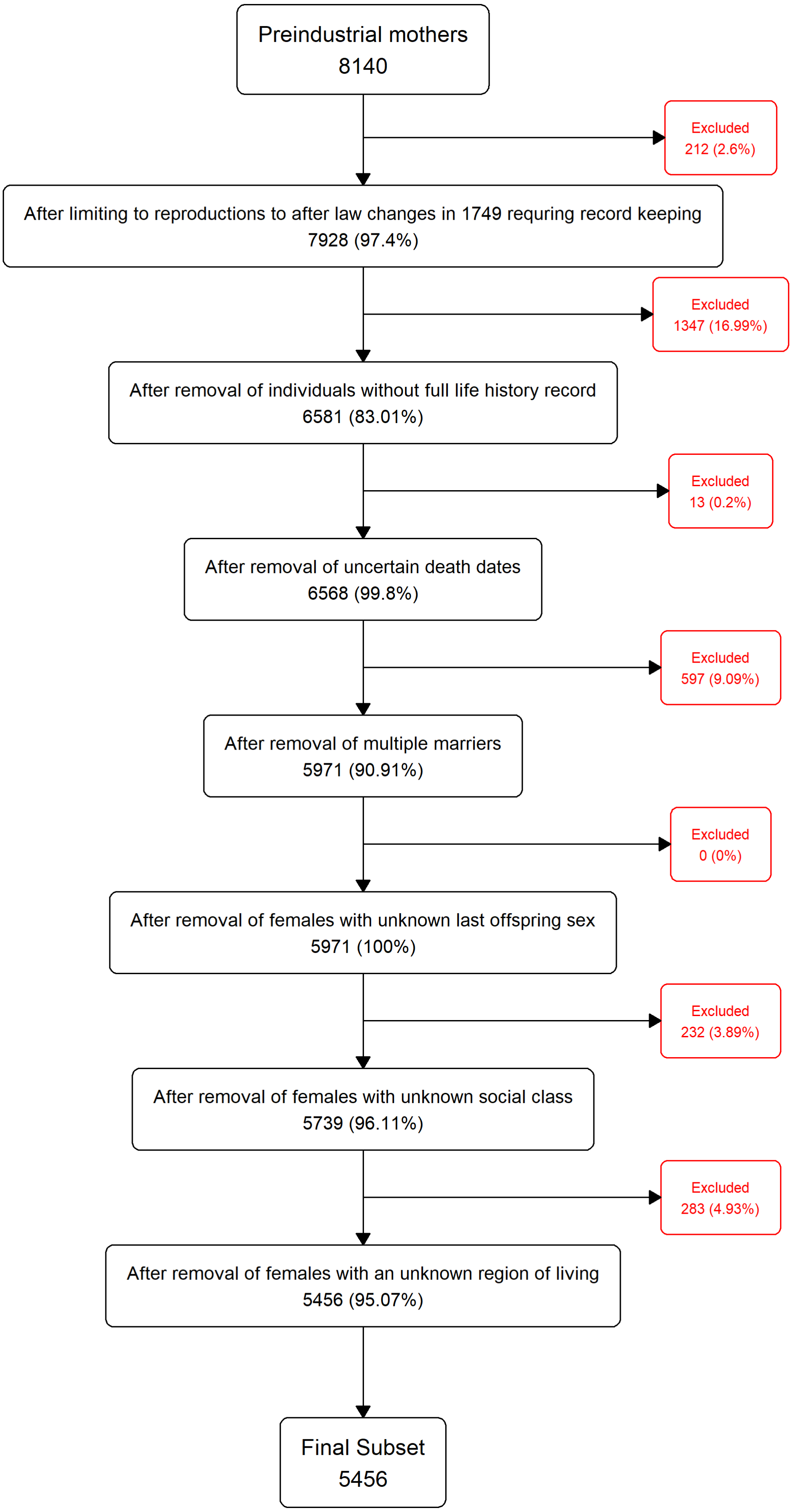


**Figure S1:** Flowchart showing data selection of mothers for the full dataset (N = 5,456).

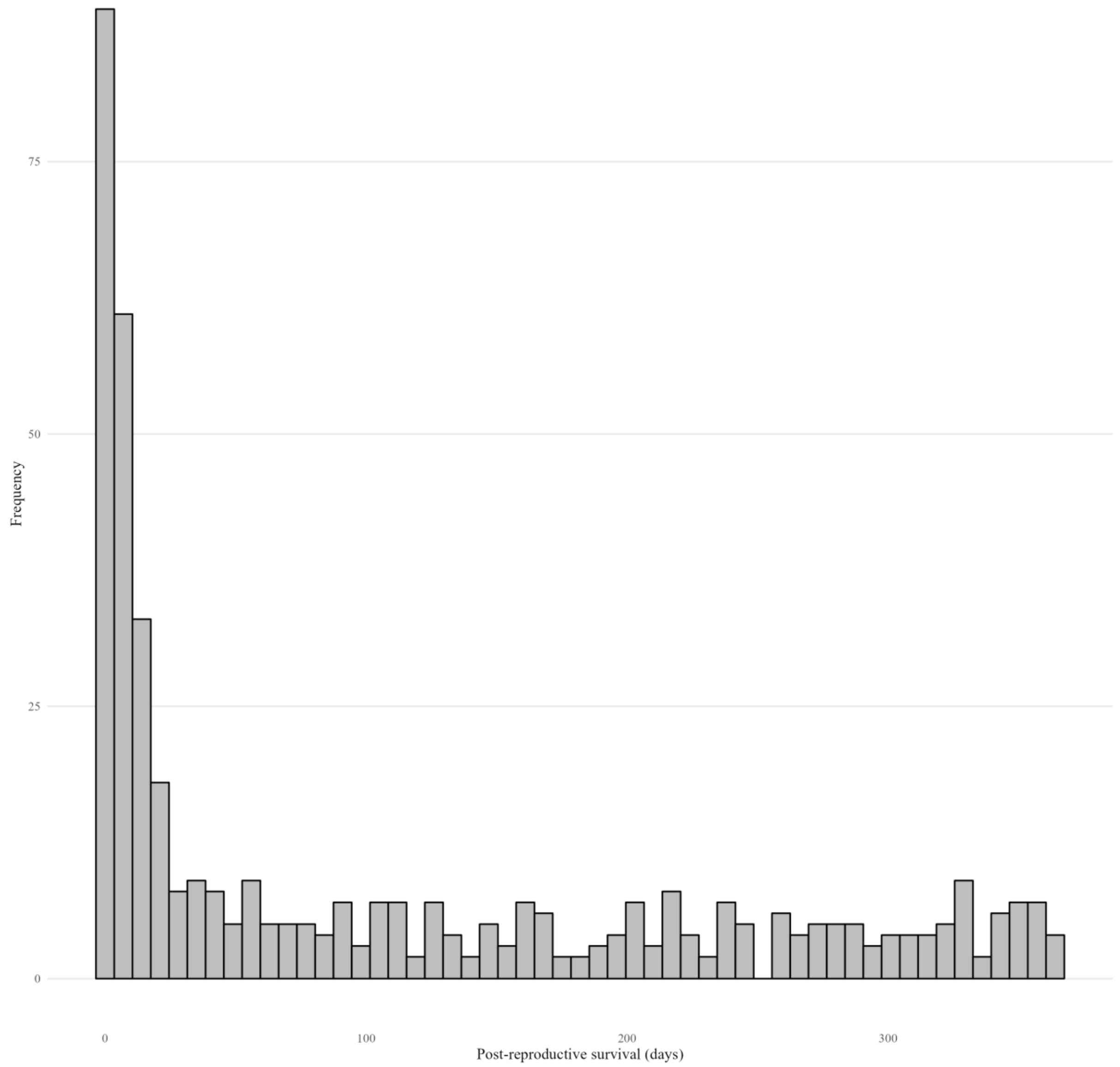


Figure S2: Histogram of deaths post-reproduction in days for mothers dying less than one-year after their last birth (n = 446). Each bar represents one week.

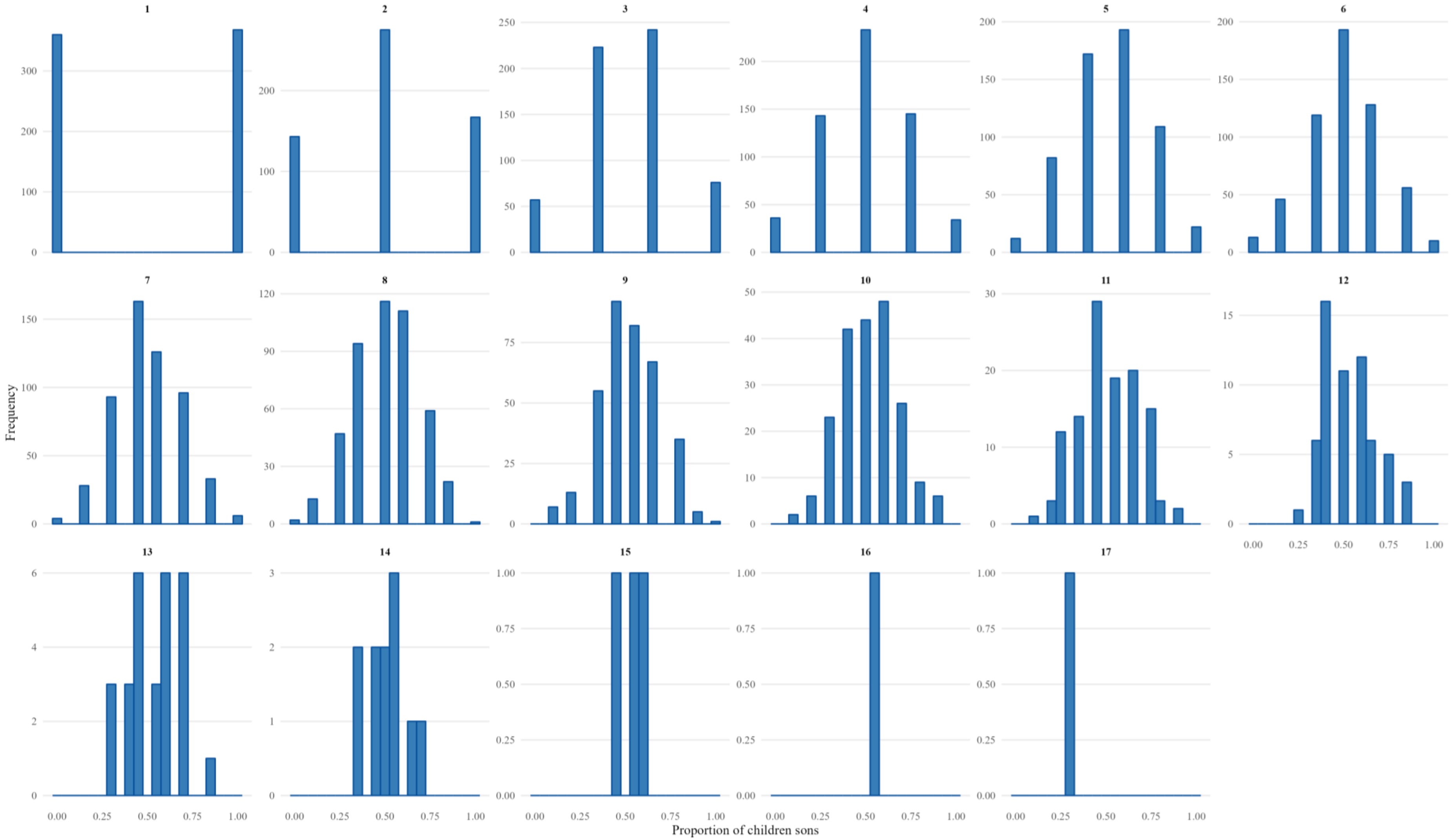


**Figure S3:** Histograms showing the proportion of children who were sons grouped by the mothers' total number of children in the full dataset (1-17, N = 4,456). The number of children for each group is indicated in the title of each histogram. Each bar width is 0.05 proportion of children sons.

Table S1: Generalised linear mixed model with all non-significant interactions included, showing conditional associations between predictors and maternal survival one year after last birth using the full dataset (N = 5,456). Estimates and variances reported are the posterior distribution median and 95% credible intervals, rounded to three significant digits. Significant effects (Pd, probability of direction > 0.975) are in bold.

| Fixed effect | Estimate [95% CI] | Pd |
|---|---|---|
| Intercept | 1.3 [-1.22 – 3.86] | 0.84 |
| Age at last reproduction | -0.057 [-0.222 – 0.106] | 0.742 |
| **Age at last reproduction^2** | **0.00334 [0.000834 – 0.0059]** | **0.998** |
| Sex (male) | -0.15 [-0.426 – 0.124] | 0.859 |
| Socioeconomic status (middle) | -0.938 [-1.96 – 0.149] | 0.956 |
| Socioeconomic status (lower) | -0.259 [-1.54 – 1.04] | 0.652 |
| Number of children | -0.155 [-0.608 – 0.26] | 0.764 |
| Number of children^2 | -0.00426 [-0.0478 – 0.0432] | 0.577 |
| Proportion sons | 0.101 [-1.13 – 1.34] | 0.566 |
| **Multiple birth (yes)** | **-0.698 [-1.18 – -0.175]** | **0.995** |
| Region (eastern Finland) | -0.203 [-0.58 – 0.199] | 0.845 |
| **Region (northern Finland)** | **-0.33 [-0.596 – -0.0473]** | **0.99** |
| Region (south-west Finland) | 0.00818 [-0.264 – 0.277] | 0.519 |
| **Number of children x Socioeconomic status (middle)** | **0.621 [0.00876 – 1.23]** | **0.977** |
| Number of children x Socioeconomic status (lower) | 0.291 [-0.535 – 1.13] | 0.754 |
| Number of children^2 x Socioeconomic status (middle) | -0.0551 [-0.117 – 0.008] | 0.958 |
| Number of children^2 x Socioeconomic status (lower) | -0.0225 [-0.117 – 0.0745] | 0.67 |
| Socioeconomic status (middle) x Proportion of children who were sons | 0.0615 [-1.65 – 1.8] | 0.528 |

| | | |
|---|---|---|
| Socioeconomic status (lower) x Proportion of children who were sons | 0.776 [-1.25 – 2.87] | 0.784 |
| Number of children x Proportion of children who were sons | -0.336 [-1.12 – 0.435] | 0.807 |
| Number of children^2 x Proportion of children who were sons | 0.0528 [-0.0316 – 0.142] | 0.892 |
| Number of children x Proportion of children who were sons x Socioeconomic status (middle) | -0.267 [-1.34 – 0.852] | 0.676 |
| Number of children x Proportion of children who were sons x Socioeconomic status (lower) | -0.436 [-1.91 – 0.964] | 0.725 |
| Number of children^2 x Proportion of children who were sons x Socioeconomic status (middle) | 0.0148 [-0.106 – 0.133] | 0.597 |
| Number of children^2 x Proportion of children who were sons x Socioeconomic status (lower) | 0.0252 [-0.15 – 0.203] | 0.604 |
| **Random Effects** | **Variance** | **Levels** |
| Birth year of last child | 0.444 [0.295 – 0.615] | 138 |

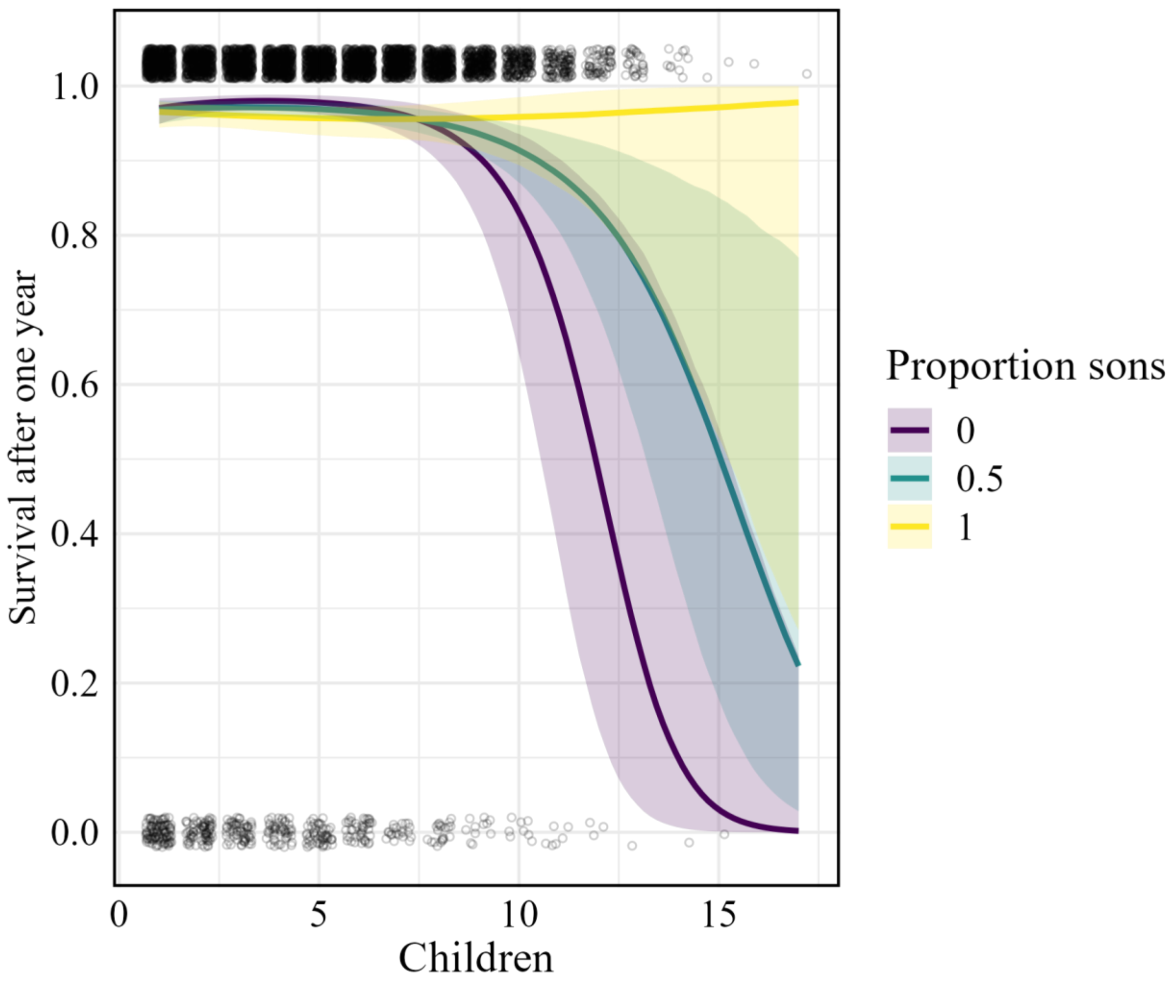


**Figure S4:** Predicted survival probabilities of mothers according to their number of children and the proportion of those children that were sons extended across all numbers of children from a generalised linear mixed model performed on the full dataset (N = 5,456). For visualisation, we predict for mothers who had only daughters (purple), an equal proportion of sons and daughters (turquoise), and only sons (yellow), across all numbers of children. Survival probabilities were predicted using posterior samples and median values or baseline values for other predictors: age at last reproduction, 38.7; Central Finland; sex of last child, male; intermediate socioeconomic status; and singleton births. Shaded areas denote 95% credible intervals. Points show surviving (above) and deceased mothers (below) are jittered for visualisation. Full model results are shown in Table 1.

Table S2: Generalised linear mixed model showing conditional associations between predictors and maternal survival one year after last birth from only mothers with 10 or fewer children matching the model structure of Table 1 (N = 5,234). Estimates and variances reported are the posterior distribution median and 95% credible intervals rounded to three significant digits. Significant effects (Pd, probability of direction > 0.975) are in bold. Model results from the models with all non-significant interactions removed and included are shown in Tables S3 and S4, respectively.

| **Fixed effect** | **Estimate [95% CI]** | **Pd** |
|---|---|---|
| Intercept | 0.361 [-2.14 – 2.9] | 0.62 |
| Age at last reproduction | 0.00219 [-0.157 – 0.162] | 0.511 |
| Age at last reproduction^2 | 0.00239 [-1.34e-05 – 0.00487] | 0.973 |
| Sex (male) | -0.133 [-0.42 – 0.144] | 0.817 |
| Number of children | -0.0924 [-0.486 – 0.312] | 0.681 |
| Number of children^2 | -0.00628 [-0.0512 – 0.0392] | 0.602 |
| Proportion of children who were sons | 0.343 [-0.514 – 1.27] | 0.775 |
| Socioeconomic status (middle) | -0.711 [-1.44 – 0.0566] | 0.966 |
| Socioeconomic status (lower) | 0.254 [-0.578 – 1.1] | 0.732 |
| **Multiple birth (yes)** | **-0.758 [-1.26 – -0.208]** | **0.997** |
| Region (eastern Finland) | -0.204 [-0.585 – 0.196] | 0.849 |
| **Region (northern Finland)** | **-0.38 [-0.656 – -0.104]** | **0.994** |
| Region (south-west Finland) | -0.0204 [-0.292 – 0.254] | 0.556 |
| Number of children x Proportion of children who were sons | -0.543 [-1.16 – 0.0447] | 0.963 |
| Number of children^2 x Proportion of children who were sons | 0.0674 [-0.00477 – 0.141] | 0.965 |
| Number of children x Socioeconomic status (middle) | 0.346 [-0.0368 – 0.718] | 0.961 |
| Number of children^2 x Socioeconomic status (middle) | -0.0312 [-0.0693 – 0.00851] | 0.94 |
| Number of children x Socioeconomic status (lower) | -0.0169 [-0.464 – 0.426] | 0.528 |
| Number of children^2 x Socioeconomic status (lower) | 0.00116 [-0.0463 – 0.0533] | 0.521 |
| **Random Effects** | **Variance** | **Levels** |
| Birth year of last child | 0.445 [0.29 – 0.613] | 138 |

Table S3: Generalised linear mixed model showing conditional associations between predictors and maternal survival one year after last birth from only mothers with 10 or fewer children with all non-significant interactions removed, except for the marginally significant interaction between number of children and proportion sons (N = 5,234). Estimates and variances reported are the posterior distribution median and 95% credible intervals rounded to three significant digits. Significant effects (Pd, probability of direction > 0.975) are in bold. Model results with all non-significant interactions included are shown in Table S4.

| Fixed effect | Estimate [95% CI] | Pd |
|---|---|---|
| Intercept | 0.228 [-2.26 – 2.65] | 0.569 |
| Age at last reproduction | -0.00124 [-0.162 – 0.159] | 0.507 |
| Age at last reproduction^2 | 0.00245 [-2.16e-05 – 0.00495] | 0.974 |
| Sex (male) | -0.142 [-0.415 – 0.135] | 0.845 |
| Number of children | 0.0221 [-0.326 – 0.376] | 0.552 |
| Number of children^2 | -0.0168 [-0.0581 – 0.0229] | 0.795 |
| Proportion of children who were sons | 0.351 [-0.503 – 1.26] | 0.785 |
| Socioeconomic status (middle) | -0.0308 [-0.266 – 0.229] | 0.592 |
| Socioeconomic status (lower) | 0.236 [-0.0519 – 0.522] | 0.951 |
| **Multiple birth (yes)** | **-0.757 [-1.24 – -0.206]** | **0.998** |
| Region (eastern Finland) | -0.219 [-0.599 – 0.169] | 0.869 |
| **Region (northern Finland)** | **-0.401 [-0.666 – -0.108]** | **0.999** |
| Region (south-west Finland) | -0.0287 [-0.293 – 0.238] | 0.581 |
| Number of children x Proportion of children who were sons | -0.547 [-1.15 – 0.0294] | 0.969 |
| Number of children^2 x Proportion of children who were sons | 0.0677 [-0.0019 – 0.142] | 0.973 |
| **Random Effects** | **Variance** | **Levels** |
| Birth year of last child | 0.434 [0.276 – 0.6] | 138 |

Table S4: Generalised linear mixed model showing conditional associations between predictors and maternal survival one year after last birth from only mothers with 10 or fewer children with non-significant interactions included (N = 5,234). Estimates and variances reported are the posterior distribution median and 95% credible intervals rounded to three significant digits. Significant effects (Pd, probability of direction > 0.975) are in bold.

| **Fixed effect** | **Estimate [95% CI]** | **Pd** |
|---|---|---|
| Intercept | 0.445 [-2.13 – 3.14] | 0.634 |
| Age at last reproduction | 0.00543 [-0.163 – 0.169] | 0.523 |
| Age at last reproduction^2 | 0.00234 [-0.000123 – 0.00497] | 0.969 |
| Sex (male) | -0.137 [-0.417 – 0.138] | 0.841 |
| Socioeconomic status (middle) | -0.778 [-1.97 – 0.443] | 0.887 |
| Socioeconomic status (lower) | 0.163 [-1.25 – 1.57] | 0.593 |
| Number of children | -0.17 [-0.677 – 0.324] | 0.75 |
| Number of children^2 | -0.00218 [-0.0567 – 0.056] | 0.529 |
| Proportion of children who were sons | 0.179 [-1.13 – 1.48] | 0.609 |
| **Multiple birth (yes)** | **-0.743 [-1.26 – -0.225]** | **0.997** |
| Region (eastern Finland) | -0.216 [-0.58 – 0.183] | 0.864 |
| **Region (northern Finland)** | **-0.39 [-0.67 – -0.117]** | **0.997** |
| Region (south-west Finland) | -0.0174 [-0.3 – 0.256] | 0.553 |
| Number of children x Socioeconomic status (middle) | 0.521 [-0.223 – 1.3] | 0.907 |
| Number of children x Socioeconomic status (lower) | -0.0913 [-1.03 – 0.837] | 0.583 |
| Number of children^2 x Socioeconomic status (middle) | -0.0465 [-0.135 – 0.043] | 0.849 |
| Number of children^2 x Socioeconomic status (lower) | 0.0352 [-0.0781 – 0.154] | 0.718 |
| Socioeconomic status (middle) x Proportion of children who were sons | 0.154 [-1.78 – 2.05] | 0.558 |
| Socioeconomic status (lower) x Proportion of children who were sons | 0.211 [-1.94 – 2.41] | 0.575 |

| | | |
|---|---|---|
| Number of children x Proportion of children who were sons | -0.393 [-1.26 – 0.503] | 0.804 |
| Number of children^2 x Proportion of children who were sons | 0.0598 [-0.0446 – 0.162] | 0.875 |
| Number of children x Proportion of children who were sons x Socioeconomic status (middle) | -0.351 [-1.65 – 0.965] | 0.692 |
| Number of children x Proportion of children who were sons x Socioeconomic status (lower) | 0.122 [-1.41 – 1.64] | 0.56 |
| Number of children^2 x Proportion of children who were sons x Socioeconomic status (middle) | 0.0288 [-0.132 – 0.188] | 0.634 |
| Number of children^2 x Proportion of children who were sons x Socioeconomic status (lower) | -0.0618 [-0.253 – 0.131] | 0.733 |
| **Random Effects** | **Variance** | **Levels** |
| Birth year of last child | 0.439 [0.28 – 0.612] | 138 |

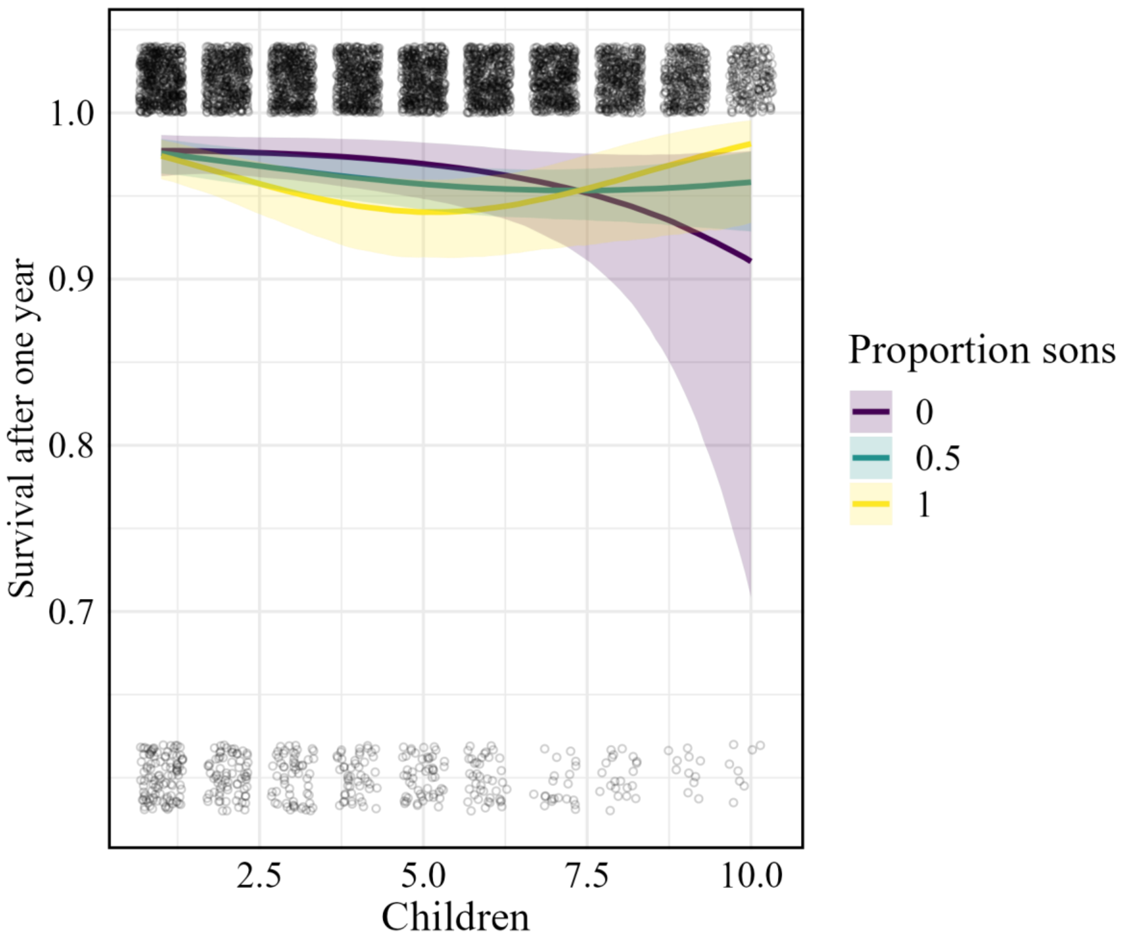


**Figure S5:** Predicted survival probabilities of mothers according to their number of children and the proportion of those children that were sons extended across all numbers of children from a generalised linear mixed model performed on mothers with 10 or fewer children (N = 5,234). For visualisation, we predict for mothers who had only daughters (purple), an equal proportion of sons and daughters (turquoise), and only sons (yellow) and across all numbers of children. Survival probabilities were predicted using posterior samples and median values or baseline values for other predictors: age at last reproduction, 38.4; Central Finland; sex of last child, male; intermediate socioeconomic status; and singleton births. Shaded areas denote 95% credible intervals. Points show surviving (above) and deceased mothers (below) are jittered for visualisation. Full model results are shown in Table S3.

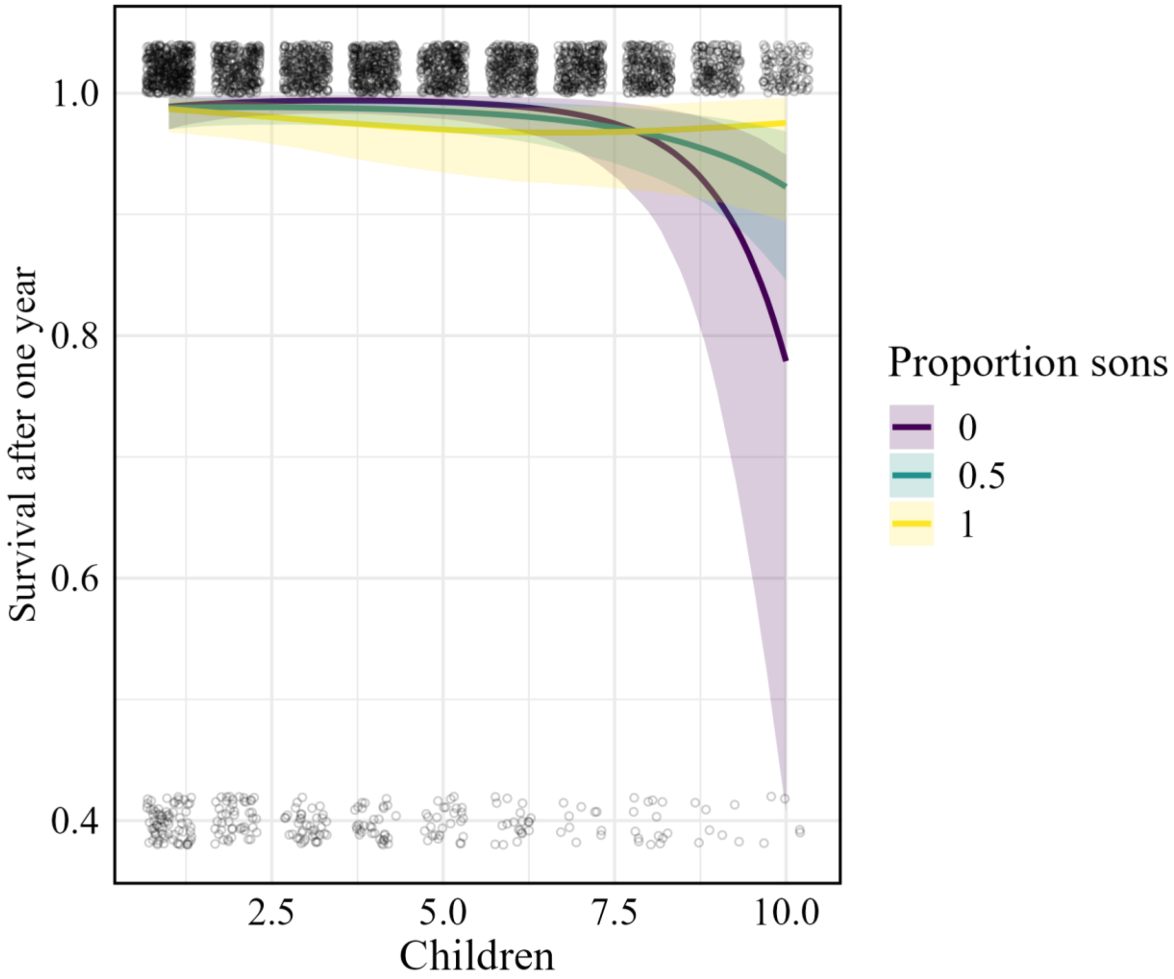


**Figure S6:** Predicted survival probabilities of mothers according to their number of children and the proportion of those children that were sons from a generalised linear mixed model performed on a reduced dataset accounting for familial differences (N = 3,246). For visualisation, we predict for mothers who had only daughters (purple), an equal proportion of sons and daughters (turquoise), and only sons (yellow) and across the 5$^{th}$ to 95$^{th}$ percentiles for number of children (1-10). Predicted survival probabilities extended across all numbers of children are plotted in Figure SX. Survival probabilities were predicted using posterior samples and median values or baseline values for other predictors: age at last reproduction, 38.4; Central Finland; sex of last child, male; intermediate socioeconomic status; and singleton births. Shaded areas denote 95% credible intervals. Points show surviving (above) and deceased mothers (below) are jittered for visualisation. Full model results are shown in Table S5.

Table S5: Generalised linear mixed model showing conditional associations between predictors and maternal survival one year after last birth accounting for familial environment (N = 3,246). Estimates and variances reported are the posterior distribution median and 95% credible intervals rounded to three significant digits. Significant effects (Pd, probability of direction > 0.975) are in bold. Model results with non-significant interactions included are shown in Table S6.

| **Fixed effect** | **Estimate [95% CI]** | **Pd** |
|---|---|---|
| Intercept | 1.01 [-2.79 – 4.92] | 0.699 |
| Age at last reproduction | -0.0488 [-0.293 – 0.201] | 0.654 |
| **Age at last reproduction^2** | **0.0041 [0.000334 – 0.008]** | **0.984** |
| Sex (male) | -0.321 [-0.762 – 0.102] | 0.93 |
| Number of children | 0.0725 [-0.413 – 0.535] | 0.618 |
| Number of children^2 | -0.033 [-0.0793 – 0.0163] | 0.903 |
| Proportion sons | 0.765 [-0.449 – 1.94] | 0.898 |
| **Socioeconomic status (middle)** | **-1.3 [-2.37 – -0.323]** | **0.996** |
| Socioeconomic status (lower) | -0.169 [-1.25 – 0.901] | 0.625 |
| **Multiple birth (yes)** | **-0.954 [-1.85 – -0.0783]** | **0.984** |
| Region (eastern Finland) | -0.312 [-0.936 – 0.314] | 0.833 |
| Region (northern Finland) | -0.357 [-0.789 – 0.0571] | 0.953 |
| Region (south-west Finland) | -0.00564 [-0.439 – 0.448] | 0.514 |
| **Number of children x Proportion of children who were sons** | **-1.05 [-1.86 – -0.284]** | **0.997** |
| **Number of children^2 x Proportion of children who were sons** | **0.122 [0.04 – 0.215]** | **0.999** |
| **Number of children x Socioeconomic status (middle)** | **0.592 [0.164 – 1.06]** | **0.998** |
| **Number of children^2 x Socioeconomic status (middle)** | **-0.06 [-0.103 – -0.0228]** | **0.999** |
| Number of children x Socioeconomic status (lower) | 0.26 [-0.284 – 0.826] | 0.821 |
| Number of children^2 x Socioeconomic status (lower) | -0.0354 [-0.094 – 0.0217] | 0.891 |
| **Random Effects** | **Variance** | **Levels** |
| Family | 1.27 [0.439 – 1.98] | 2,199 |

| | | |
|---|---|---|
| Birth year of last child | 0.61 [0.363 – 0.911] | 136 |

Table S6: Generalised linear mixed model showing conditional associations between predictors and maternal survival one year after last birth accounting for familial environment with non-significant interactions included (N = 3,246). Estimates and variances reported are the posterior distribution median and 95% credible intervals rounded to three significant digits. Significant effects (Pd, probability of direction > 0.975) are in bold.

| Fixed effect | Estimate [95% CI] | Pd |
|---|---|---|
| Intercept | 0.856 [-3.48 – 5.14] | 0.645 |
| Age at last reproduction | -0.0351 [-0.302 – 0.227] | 0.601 |
| **Age at last reproduction^2** | **0.00402 [0.000125 – 0.00823]** | **0.977** |
| Sex (male) | -0.343 [-0.823 – 0.0915] | 0.939 |
| Socioeconomic status (middle) | -1.05 [-2.8 – 0.632] | 0.893 |
| Socioeconomic status (lower) | -0.531 [-2.46 – 1.31] | 0.719 |
| Number of children | 0.0318 [-0.651 – 0.647] | 0.537 |
| Number of children^2 | -0.0384 [-0.104 – 0.0321] | 0.86 |
| Proportion sons | 0.931 [-1.07 – 2.92] | 0.823 |
| **Multiple birth (yes)** | **-1.03 [-1.98 – -0.103]** | **0.984** |
| Region (eastern Finland) | -0.329 [-1.03 – 0.358] | 0.829 |
| Region (northern Finland) | -0.378 [-0.852 – 0.0509] | 0.955 |
| Region (south-west Finland) | -0.0151 [-0.447 – 0.46] | 0.524 |
| Number of children x Socioeconomic status (middle) | 0.608 [-0.351 – 1.63] | 0.897 |
| Number of children x Socioeconomic status (lower) | 0.556 [-0.644 – 1.81] | 0.809 |
| Number of children^2 x Socioeconomic status (middle) | -0.051 [-0.158 – 0.0472] | 0.856 |
| Number of children^2 x Socioeconomic status (lower) | -0.0443 [-0.179 – 0.0938] | 0.733 |
| Socioeconomic status (middle) x Proportion of children who were sons | -0.624 [-3.44 – 2.19] | 0.687 |

| | | |
|---|---|---|
| Socioeconomic status (lower) x Proportion of children who were sons | 0.536 [-2.43 – 3.6] | 0.648 |
| Number of children x Proportion of children who were sons | -1.08 [-2.36 – 0.196] | 0.953 |
| **Number of children^2 x Proportion of children who were sons** | **0.144 [0.000407 – 0.287]** | **0.976** |
| Number of children x Proportion of children who were sons x Socioeconomic status (middle) | 0.0641 [-1.79 – 1.85] | 0.527 |
| Number of children x Proportion of children who were sons x Socioeconomic status (lower) | -0.423 [-2.62 – 1.72] | 0.654 |
| Number of children^2 x Proportion of children who were sons x Socioeconomic status (middle) | -0.0296 [-0.221 – 0.169] | 0.623 |
| Number of children^2 x Proportion of children who were sons x Socioeconomic status (lower) | -0.000551 [-0.26 – 0.262] | 0.502 |
| **Random Effects** | **Variance** | **Levels** |
| Family | 1.43 [0.623 – 2.23] | 2,199 |
| Birth year of last child | 0.631 [0.375 – 0.951] | 136 |

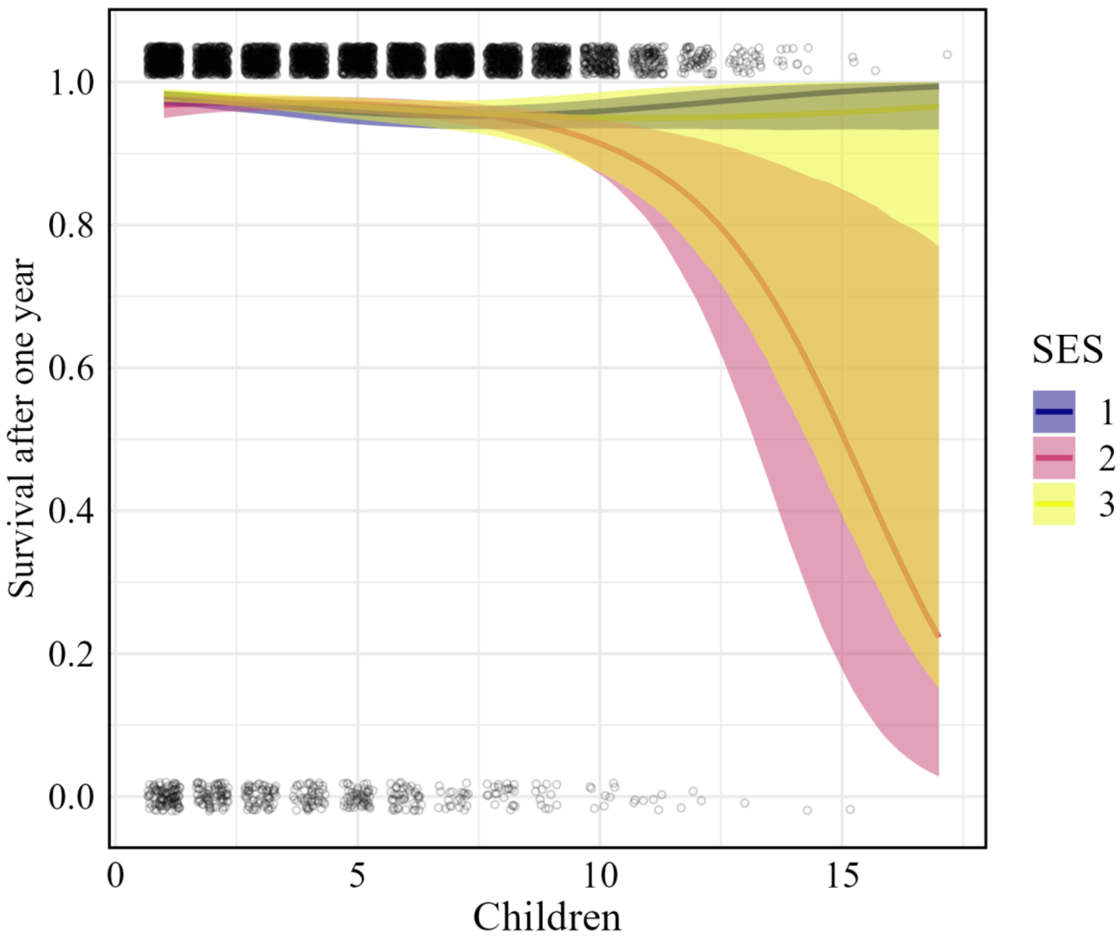


**Figure S7:** Predicted survival probabilities of mothers according to their number of children and socioeconomic status (1 = upper, purple; 2 = intermediate, pink; and 3 = lower, yellow) from a generalised linear mixed model performed on the full dataset (N = 5,456). Survival probabilities were predicted using posterior samples and median values or baseline values for other predictors: proportion of children who were sons, 0.5; age at last reproduction, 38.7; Central Finland; sex of last child, male; and singleton births. Shaded areas denote 95% credible intervals. Points show surviving (above) and deceased mothers (below) are jittered for visualisation. Full model results are shown in Table 1.

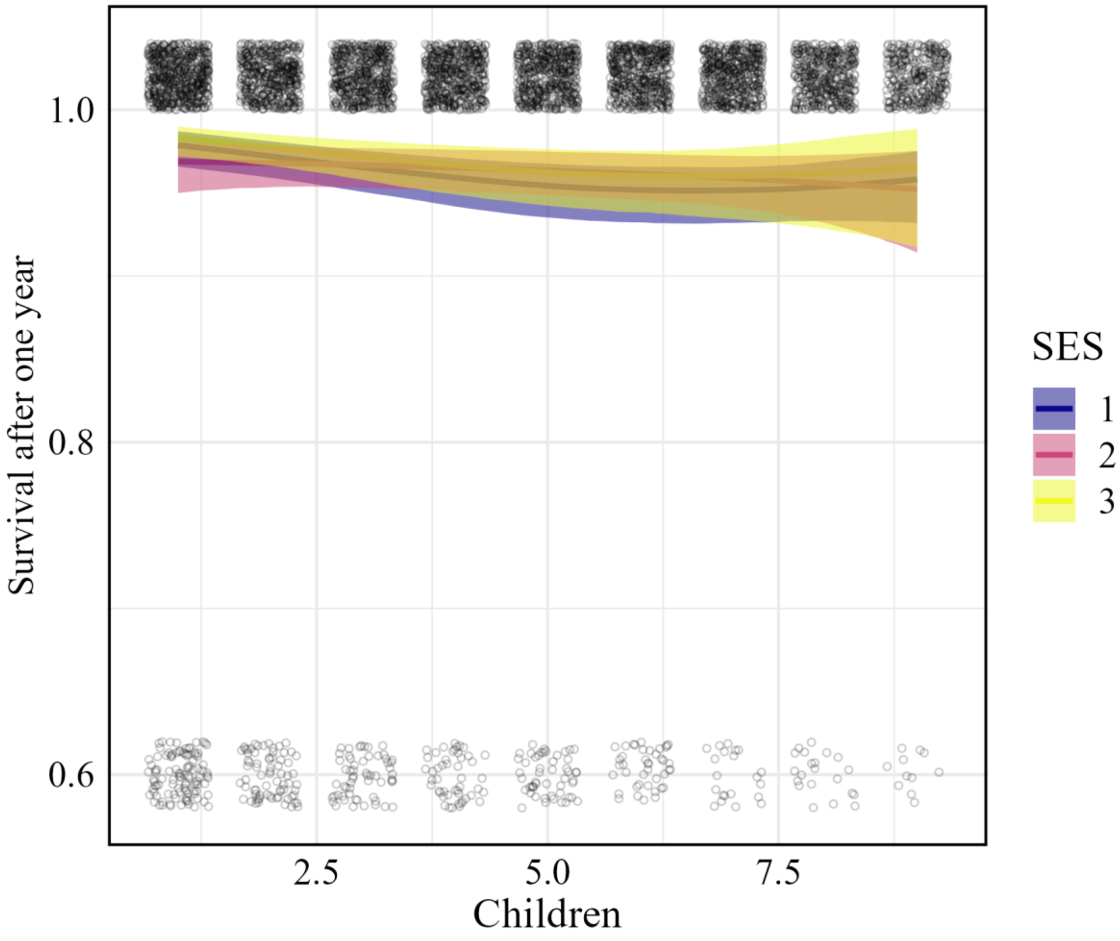


**Figure S8:** Predicted survival probabilities of mothers according to their number of children and socioeconomic status (1 = upper, purple; 2 = intermediate, pink; and 3 = lower, yellow) from a generalised linear mixed model performed on mothers with 10 or fewer children (N = 5,234). Survival probabilities were predicted using posterior samples and median values or baseline values for other predictors: proportion of children who were sons, 0.5; age at last reproduction, 38.4; Central Finland; sex of last child, male; and singleton births. Shaded areas denote 95% credible intervals. Points show surviving (above) and deceased mothers (below) are jittered for visualisation. Full model results are shown in Table S3.

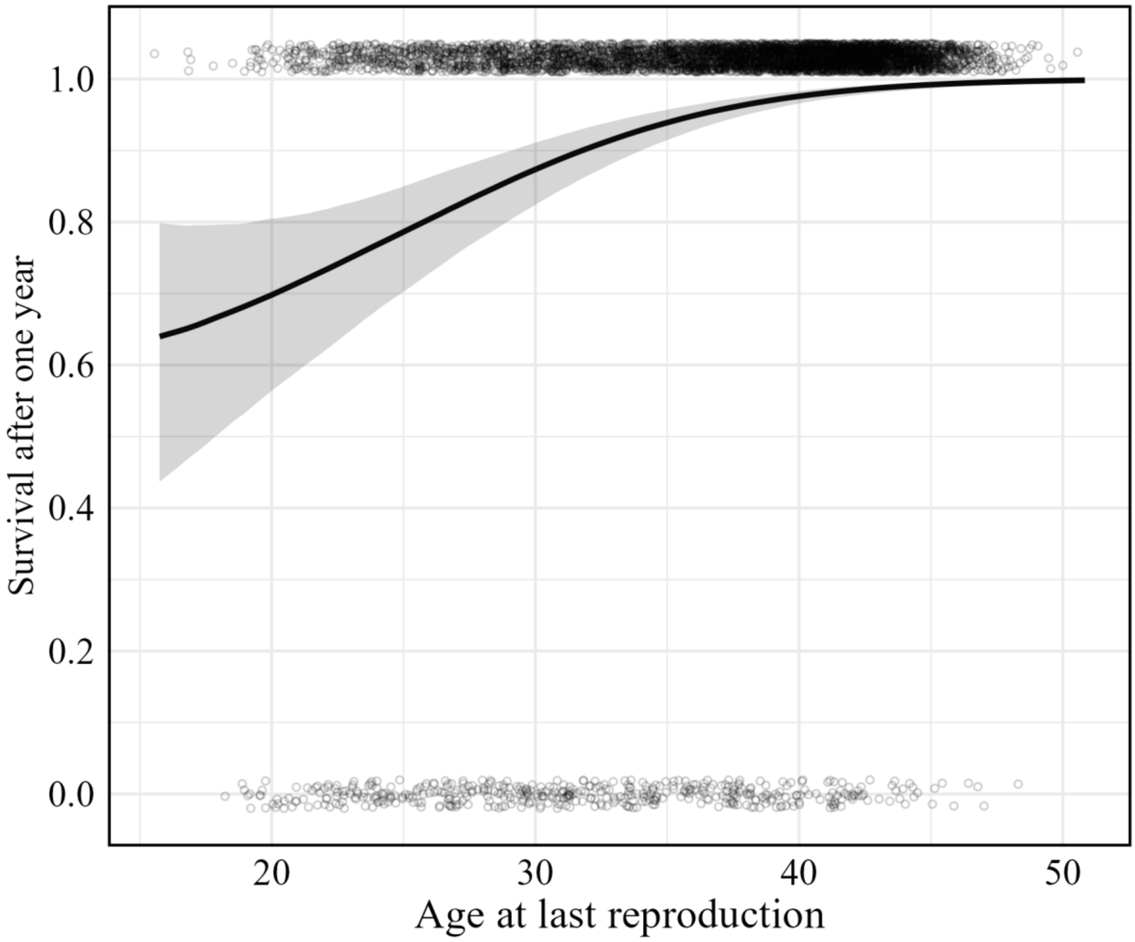


Figure S9: Survival for mothers across ages of last reproduction showing quadratic effect predicted from a generalised linear mixed model on the full dataset (N = 5,456). Survival probabilities were predicted using posterior samples and median values or baseline predictors: number of children, 5; proportion of children who were sons, 0.5; Central Finland; sex of last child, male; intermediate socioeconomic status; singleton births. Shaded areas denote 95% credible intervals. Points show surviving (above) and deceased mothers (below) are jittered for visualisation. Full model results are shown in Table 1.
